## Supplementary Information for "Innate Behavior Sequence Progression by Peptide-Mediated Interorgan Crosstalk"

**This file contains:**

Supplementary Fig. 1 – 12

Supplementary Video Legends 1-10.

### Supplementary Fig. 1.

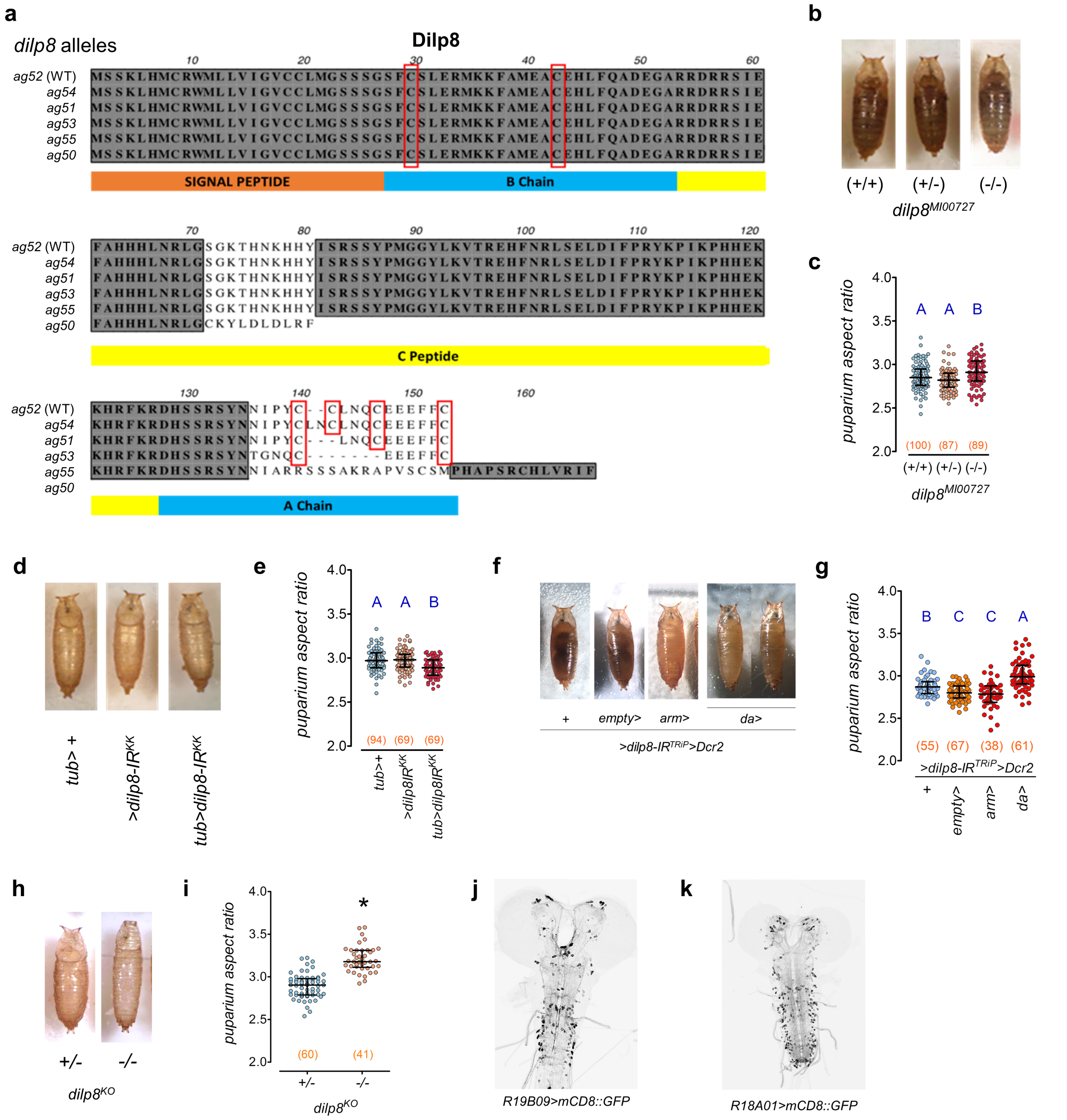


**Supplementary Fig. S1: CRISPR-Cas9-generated *dilp8* mutants and phenotypes of hypomorphs and a knockout.**

**a** Protein sequence of one wild type *dilp8*$^{ag52}$ and five *dilp8* mutant alleles (*dilp8*$^{ag50\text{-}55}$) generated by CRISPR-Cas9 mediated germline-mutagenesis directed to the 3' end that encodes the A-chain with the highly conserved cysteines essential for Dilp8 activity (highlighted with red boxes). **b, d, f, h** Photos of puparia of **b** animals carrying the hypomorphic allele *dilp8*$^{MI00727}$ in homozygosis (-/-) or heterozygosis (+/-) [(+/+), WT controls], **d** of animals ubiquitously-expressing RNAi against *dilp8* (*tub>dilp8-IR*$^{KK}$) and controls (*tub> and UAS-dilp8-IR*$^{KK}$*),* **f** of animals ubiquitously expressing *UAS-Dicer2* and

RNAi against *dilp8* (*arm>dilp8-IR*$^{TRiP}$*>Dcr2* and *da>dilp8-IR*$^{TRiP}$*>Dcr2*) and controls and **h** *dilp8*$^{KO}$ mutants (-/-) and controls (*dilp8*$^{KO}$/+ ). **c, e, g, i** Dot plots of puparium aspect ratio (AR). Dots, one animal. **b, c** *dilp8*$^{MI00727}$ mutants and **d, e** *Tub>dilp8-IR*$^{KK}$ animals do not show clear alterations in puparium shape nor a strong and consistent change in puparium AR. **f, g** Knockdown of *dilp8* with a second RNAi line alters puparium shape and aspect ratio when driven by *da>* but not by *arm>*. **h, i** *dilp8*$^{KO}$ mutants have increased puparium AR. **j, k** FlyLight[53] confocal projections of *R19B09>* and *R18A01>* driving expression of *UAS-mCD8::GFP*. Statistics: **c, e, g, i** Horizontal bar, median. Error bars: 25-75% percentiles. **c, g** Dunn's test. **d** Student-Newman-Keuls test. **h** Student's t-test . Same blue letters, $P>0.05$. **g** * $P= 6.31 E\text{-}18$. (N) number of animals (red).

**Supplementary Fig. 2.**

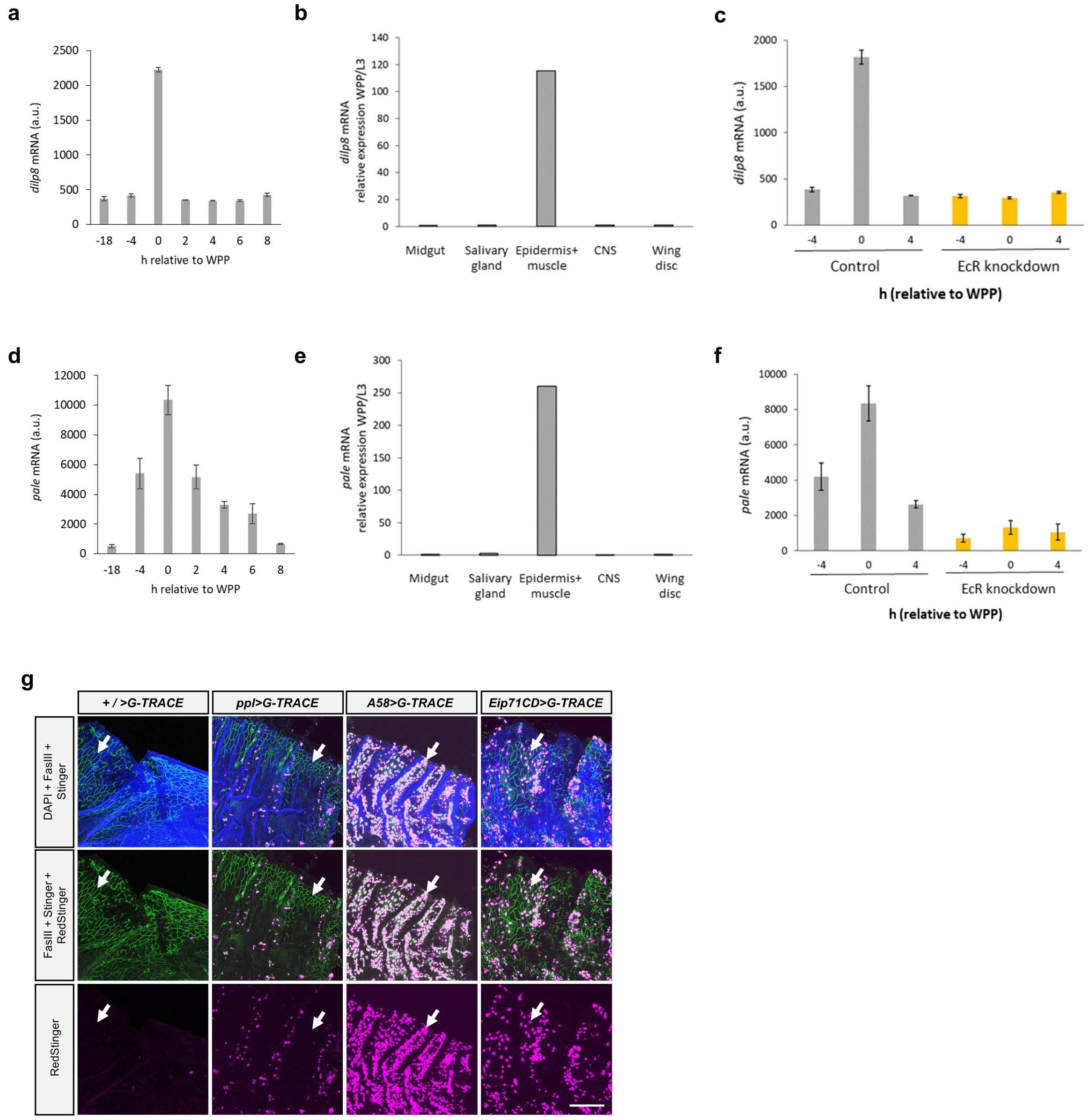


**Supplementary Fig. 2: *dilp8* and *pale* expression profiles and G-TRACE labelling of epidermal cells.**

**a-c** Expression of *dilp8* and **d-f** *pale* at pupariation (**a, b, d, e**[55]; **c, f**[56]). **a** *dilp8* and **d** *pale* expression peak with different dynamics in white prepupa (WPP T0). **b** *dilp8* and **d** *pale* are upregulated in the WPP-T0 carcass (muscle and epidermal cells). **c, f** Ecdysone receptor (EcR) knockdown prevents **c** *dilp8* and **f** *pale* upregulation at WPP T0. **g** Confocal Z-projections of immunofluorescence images of epidermal cells of the WPP T0 cuticle stained with anti-Fasciclin (FasIII; cell membranes, green) and the G-TRACE system to label past (Stinger=EGFP::NLS, nuclear green) or current (RedStinger=DsRed::NLS, nuclear magenta) expression of *ppl>, A58>*, and Eip71CD> GAL4 drivers. Blue, DAPI counterstain (nucleus) and cuticle autofluorescence. *A58>* and *Eip71CD>* drive current (nuclear magenta) and past (nuclear green) expression in the large nuclei of cuticle epidermal cells of WPP T0, *A58>* being the strongest GAL4 driver. *ppl>* drives no or only sporadic current or past cuticle epidermal-cell expression at this developmental stage. White arrows, regions of cuticle epidermal cells. Statistics: **a, c, d, f** Average. **a, c, d, f** Error bars, standard deviation.

**Supplementary Fig. 3.**

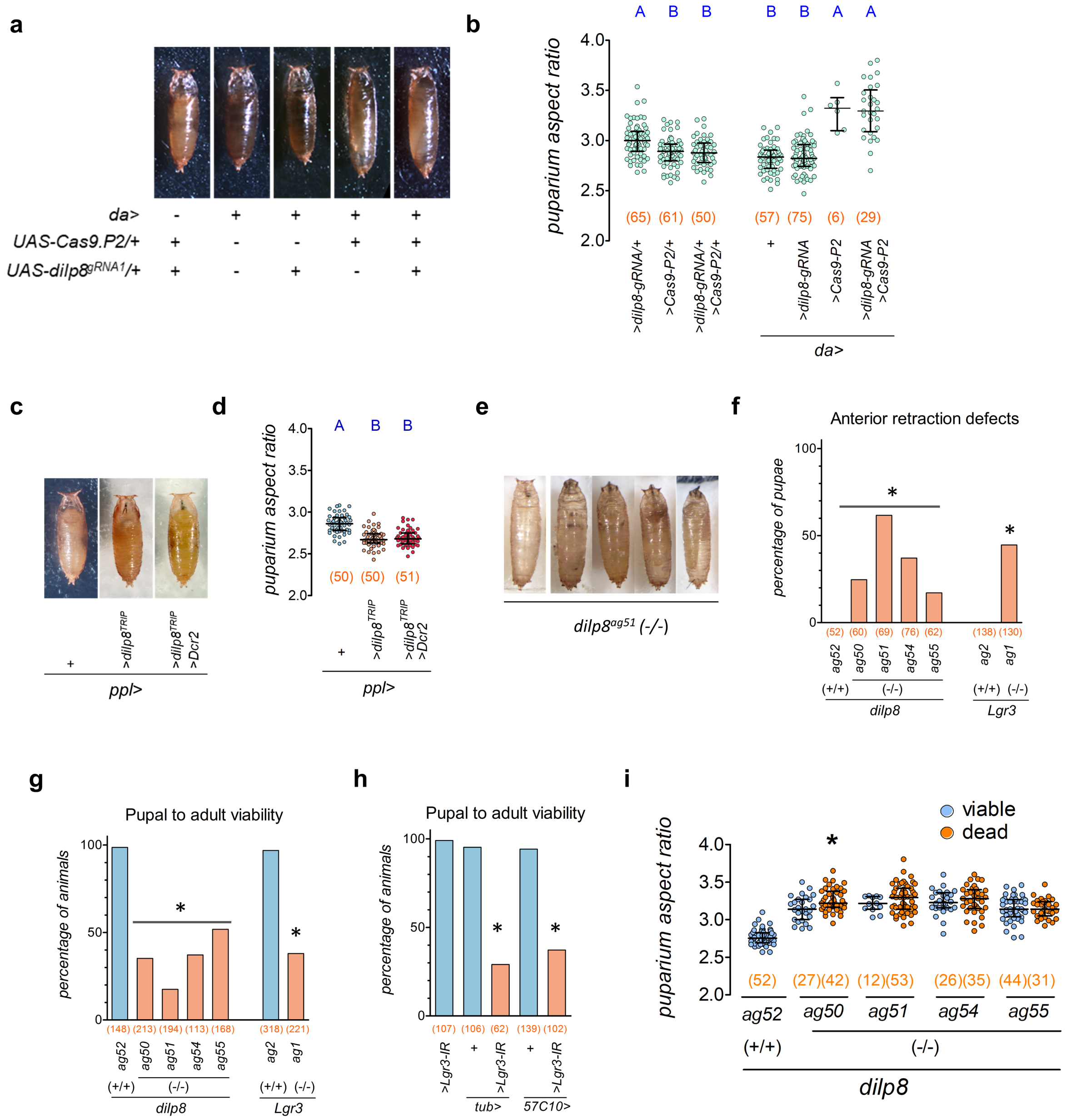


**Supplementary Fig. 3: Tissue-specific CRISPR-Cas9 and reduced pupal viability of *dilp8* and *Lgr3* loss of function.**

**a, c, e** Pictures of animals **a** ubiquitously-expressing (*da-GAL4, da>*) *UAS-Cas9P2* and/or *UAS-dilp8*$^{gRNA1}$ for targeted somatic mutagenesis of *dilp8* (Cas9 alone affects puparium morphology), **c** expressing *dilp8-IR*$^{TRIP}$ alone or combined with *Dcr2* under the fat body GAL4 driver *ppl>*, and **b** *dilp8*-mutant puparia displaying increasingly severe anterior retraction defects. **b, d, i** Dot plots of puparium aspect ratio following **b** ubiquitous *Cas9* and/or *dilp8*$^{gRNA1}$ expression, **d** knockdown of *dilp8* in the fat body, or in **i** a series of *dilp8* mutant alleles (*ag50,51,54, and* 55) or WT control (*ag52*), according to pupal viability (blue, viable; orange, dead). *ppl*>/+ animals in **c, d** are from the same batch as used in Fig. 2e, f. **f** Proportion of puparia with defective anterior retraction. **g, h.** Pupal viability is reduced in **g.** animals lacking *dilp8* and *Lgr3* or **h** following ubiquitous (*tub*>) or panneuronal (*R57C10*>) *Lgr3* knockdown. **i** Pupal viability was not associated with puparium AR, except in one of the four assayed *dilp8* mutant alleles (*ag50*). Statistics: **b, d, i** Horizontal bar, median. Error bars: 25-75% percentiles. Same blue letter, $P>0.05$. **b** Dunn’s test and **d** Student-Newman-Keuls test. **f-h** Binomial tests with Bonferroni corrections. $*P<0.001$. **i** Bonferroni’s test for multiple comparisons, $*P<0.05$ (N) number of animals (red).

**Supplementary Fig. 4.**

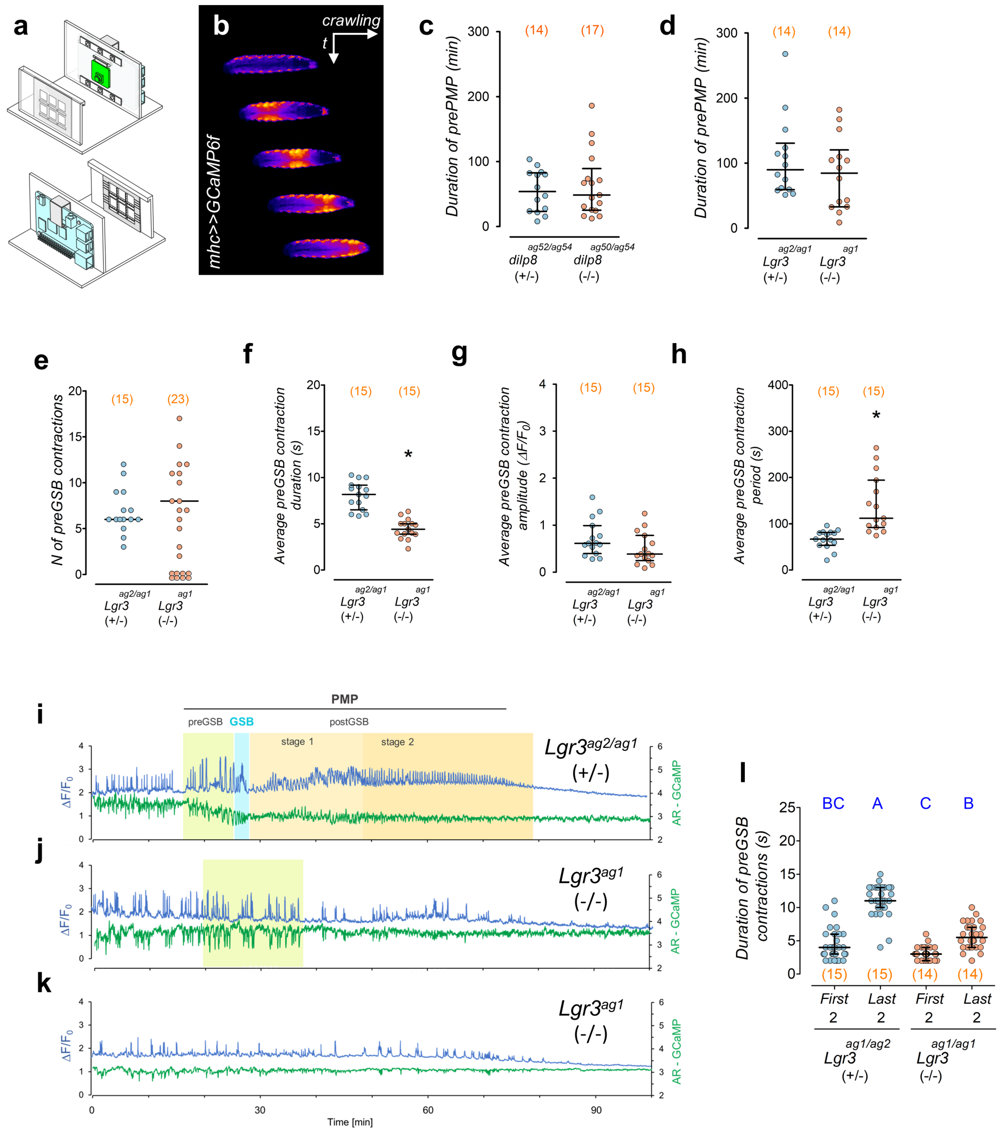


**Supplementary Fig. 4: Effects of *dilp8* and *Lgr3* on pre-PMP and pre-GSB.**

**a** Schematics of the pupariation monitoring device. **b** Time-lapse of a larva expressing the muscle-specific calcium sensor, mhc>>GCaMP, showing peristaltic contractions during larval crawling. **c-h, l** Dot plots showing **c, d** pre-pupariation motor program (pre-PMP) duration in **c** *dilp8* and **d** *Lgr3* mutants, and **d** number (dots: one animal), average **f** duration, **f** amplitude, **h** period (dots: average per larva) of pre-GSB contractions in *Lgr3* mutants and controls, or **l** duration of the first and last two pre-GSB contractions in *Lgr3* mutants and controls. **i-k** Temporal profile of GCaMP fluctuations (blue) and AR (green) in **i** control and **j, k** *Lgr3* mutant animals. **j, k** *Lgr3* mutants either show pre-GSB-like contractions (**j**), or not (**k**). **l** *Lgr3* mutants fail to increase the duration of the pre-GSB contractions with time.

Statistics: **c-h, l** Horizontal bar, median. Error bars: 25-75% percentiles. **c, d, g, h** Mann-Whitney test. **e, f** Student's t-test. **c-h** $P$=0.86, 0.26, 0.058 (animals without contractions excluded), 1.51E-08, 0.106 and <0.001, respectively. *$P$<0.001. **l** Same blue letters P>0.05, Dunn's test. (N) number of animals (red).

### Supplementary Fig. 5.

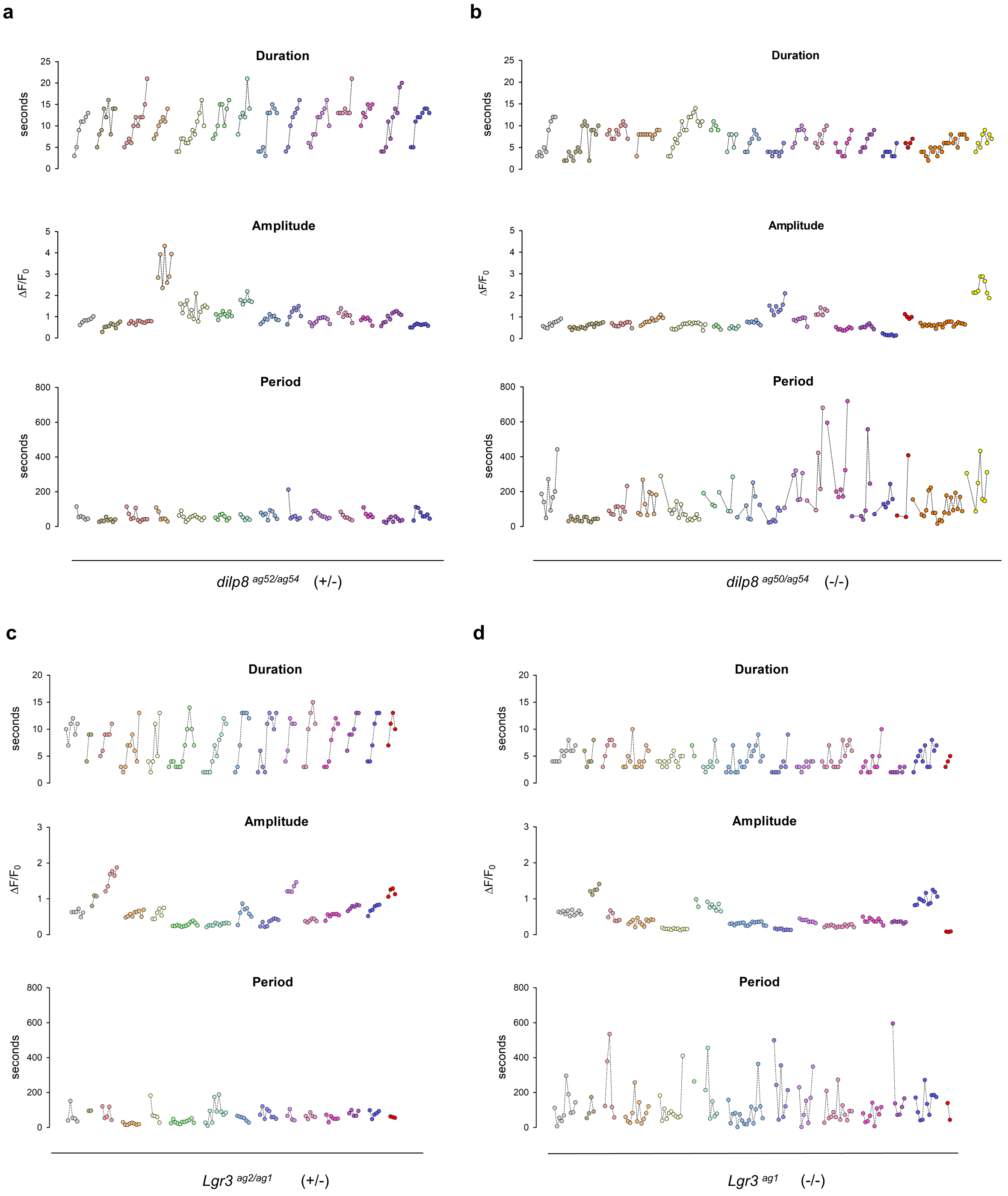


**Supplementary Fig. 5: Effects of *dilp8* and *Lgr3* on pre-GSB peak parameters.** **a-d** Duration, amplitude, and period of pre-GSB contractions (as determined by mhc>>GCaMP-fluorescence peaks) in **a, c** WT animals of the **a** *dilp8*$^{ag52/ag54}$ *(+/-)* and **c** *Lgr3*$^{ag2/ag1}$ *(+/-)* genetic background, and the **b, d** mutant: **b** *dilp8*$^{ag50/ag54}$ *(-/-)* and **d** *Lgr3*$^{ag1/ag1}$ *(-/-)*. Dots connected with a line represent consecutive contractions of one larva.

**Supplementary Fig. 6.**

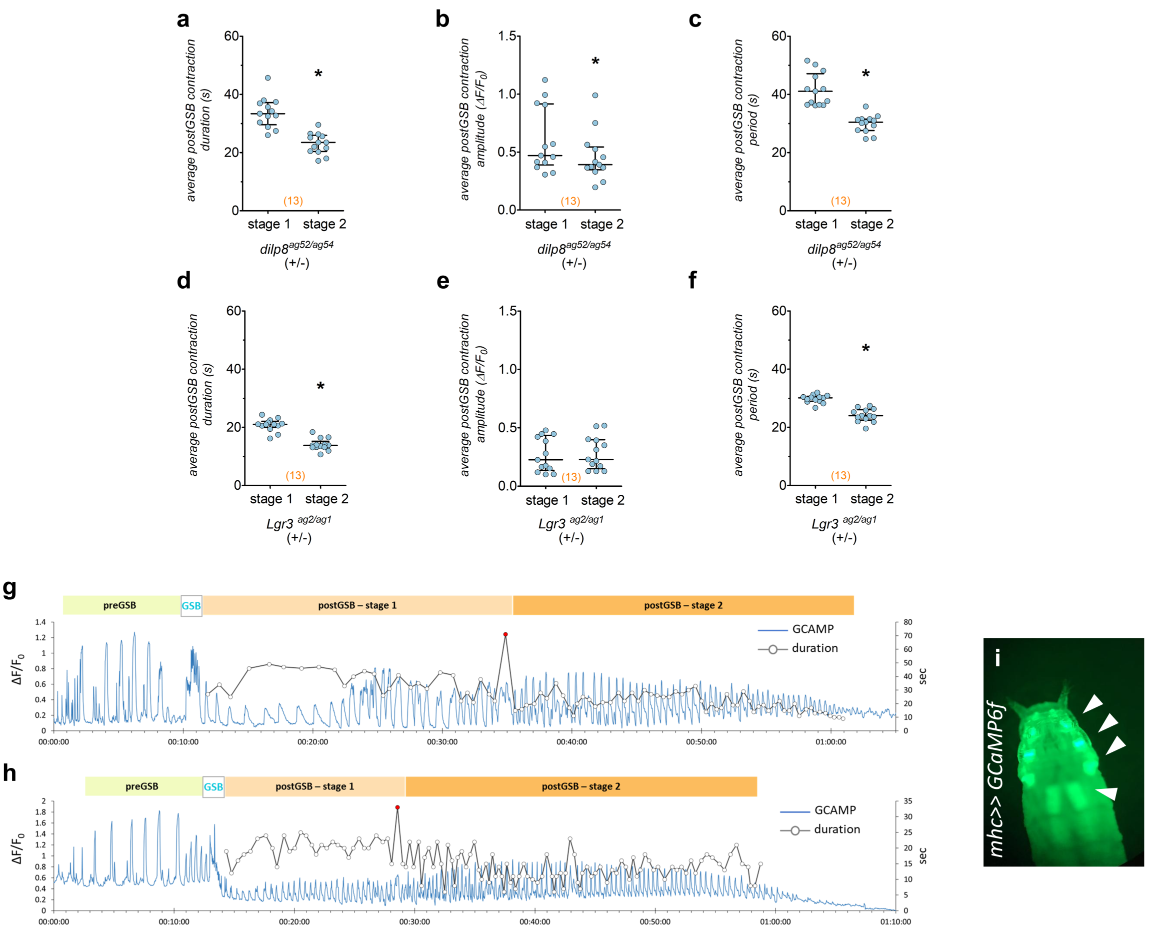


**Supplementary Fig. 6: Post-GSB stages and operculum formation program**

**a-f** Dot plots of **a, d** duration, **b, e** amplitude, and **c, f** period (dots: average per larva) of post-GSB contractions according to stages in WT animals of the **a-c** *dilp8*$^{ag52/ag54}$ *(+/-)* and **d-f** *Lgr3*$^{ag2/ag1}$ *(+/-)* genetic backgrounds. **g, h** mhc>>GCaMP-fluorescence profiles (blue line) of the pupariation motor program (PMP) of two WT animals indicating the durations of pre-GSB (light green box), GSB (white box and blue font), and post-GSB stages 1 and 2 (light and dark orange boxes, respectively). The duration of each post-GSB peak in seconds (sec) is depicted by the dark line and each white circle corresponds to one contraction. The limit between post-GSB stages 1 and 2 is marked by a contraction that is more prolonged in comparison with the neighboring contractions (marked by the red circle). **i** Dissection microscope photo taken under blue light of the mhc>>GCaMP-fluorescence pattern in a WT animal during operculum formation stage, which probably corresponds to late post-GSB-2. Statistics: **a-f** Horizontal bar, median. Error bars: 25-75% percentiles. **a, c-f** Paired Student's t test**. b** Wilcoxon signed Rank test. *$P<0.001$. (N) number of animals (red). $\Delta F/F0 = (F-F0)/F0$, being F0 the minimum value of each trace.

**Supplementary Fig. 7.**

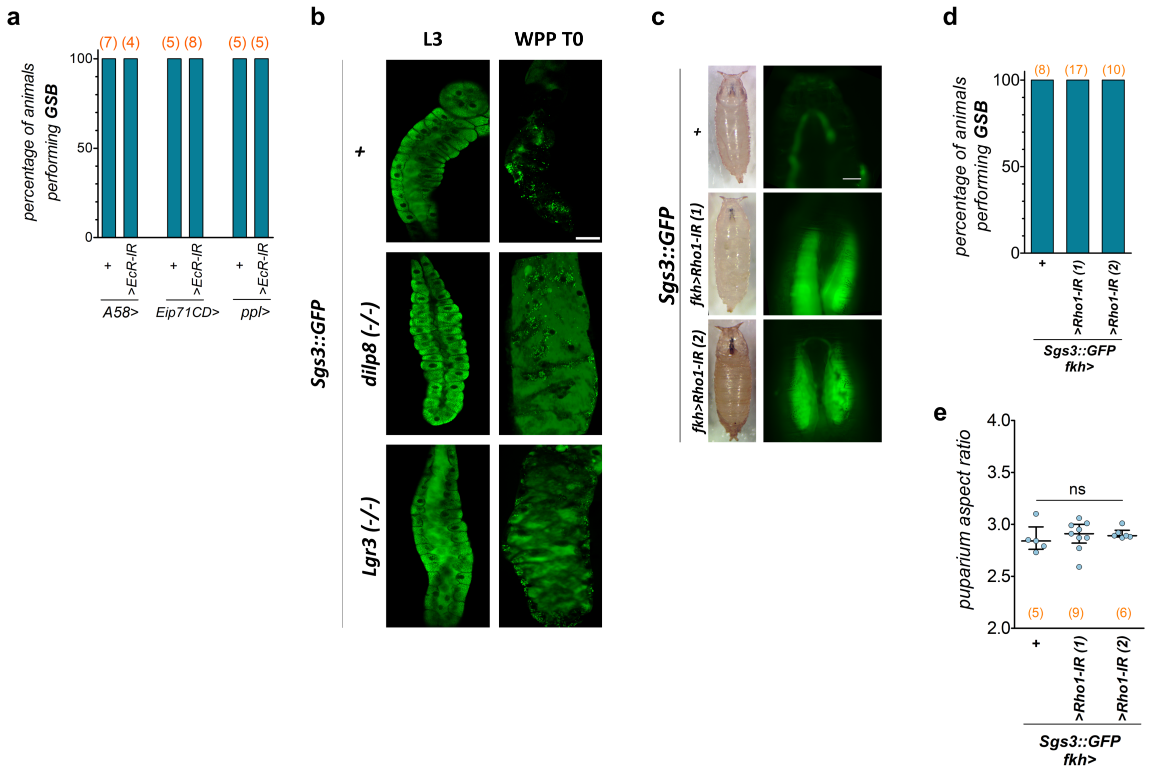


**Supplementary Fig. 7: Effects of epidermal *EcR* activity, whole-body *dilp8* and *Lgr3* activity, and glue expulsion activity on puparium morphology and GSB.**

**a, d** Percentage of animals that perform GSB upon **a** *EcR* knockdown in the epidermis (*A58>EcR-IR* and *Eip7CD>EcR-IR*) or in the fat body (*ppl>EcR-IR*), or **d** *Rho1* knockdown using two different RNAi constructs against *Rho1* (*>Rho1-IR(1)* and *>Rho1-IR(2)*) in the salivary gland (using *fkh-GAL4, fkh>*). **a** *A58>/+, Eip7CD>/+,* and *ppl>/+* controls are the same as in Fig. 5i. **b** Confocal sections of dissected salivary glands of the depicted genotypes at L3 (wandering) and WPP T0 (pupariation) stage. **c** Photos of WPP T0 puparia and Sgs3::GFP (green) showing that knockdown of *Rho1* in the salivary gland (*fkh>Rho1-IR*) impedes glue expulsion, but does not **d** affect GSB (percentage of pupa performing GSB) or **e** puparium aspect ratio (AR), depicted by dot plots. Statistics: **a, d** Binomial test with Bonferroni corrections. **e** Horizontal bar, median. Error bars: 25-75% percentiles. ANOVA. ns, not-significant, $P>0.05$. **b** *$P<0.05$. (N) number of animals (red). Scale bar: **b** 30 µm. **c** 200 µm.

**Supplementary Fig. 8.**

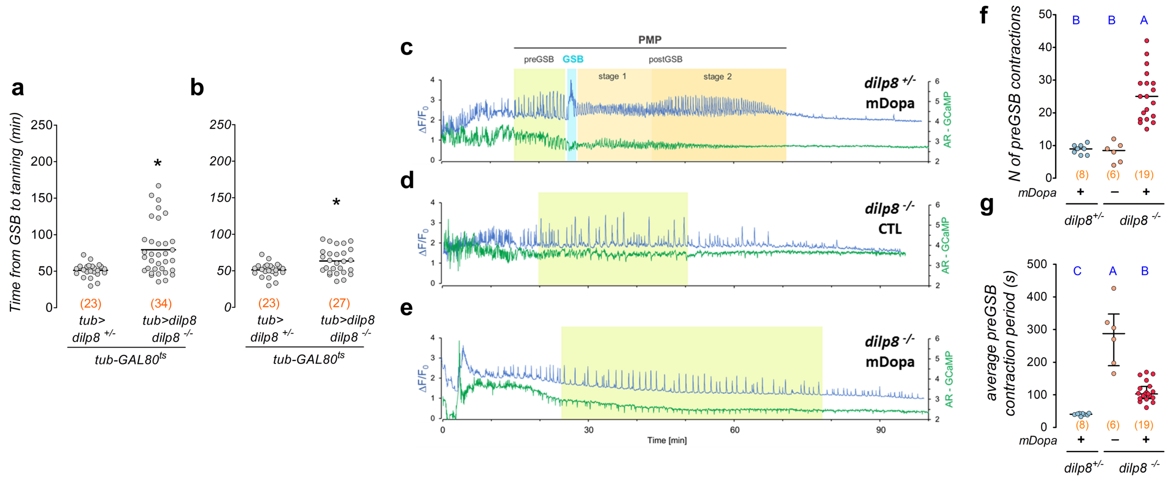


**Supplementary Fig. 8: Pupariation progression requires coupling of morphogenetic and neuromotor subprograms by the Dilp8-Lgr3 pathway.**

**a, b, f, g** Dot plots of **a, b** time from GSB to tanning following removal of animals with **a** double GSB or **b** animals with time >100 min, **f** number and **g** average period of pre-GSB contractions. **c-e** mhc>>GCaMP (blue) and aspect ratio (AR-GCaMP, green) fluctuations in **c** WT and **d, e** *dilp8* mutants treated (**c, e**) or not (**d**) with α-methyldopa. **a, b** Post- midthird instar transition-expression of *tub>dilp8* delays tanning. **c** PMP depicting stages. PMP is not overtly altered by α-methyldopa treatment of WT animals. **d** Non-treated and **d** α-methyldopa-treated *dilp8* mutants only show pre-GSB-like contractions, never transitioning to GSB. **f, g** α-methyldopa treatment **f** increases the number and decreases the average period of pre-GSB contractions in *dilp8* mutants. Statistics: Horizontal bar, **a, b** average or **F,G.** median. Error bars: 25-75%. **a, b** Mann-Whitney Rank sum test. **f, g** Dunn's test. Same blue letters, $P>0.05$. *$P<0.05$. Scale bars, 50 µm.

**Supplementary Fig. 9.**

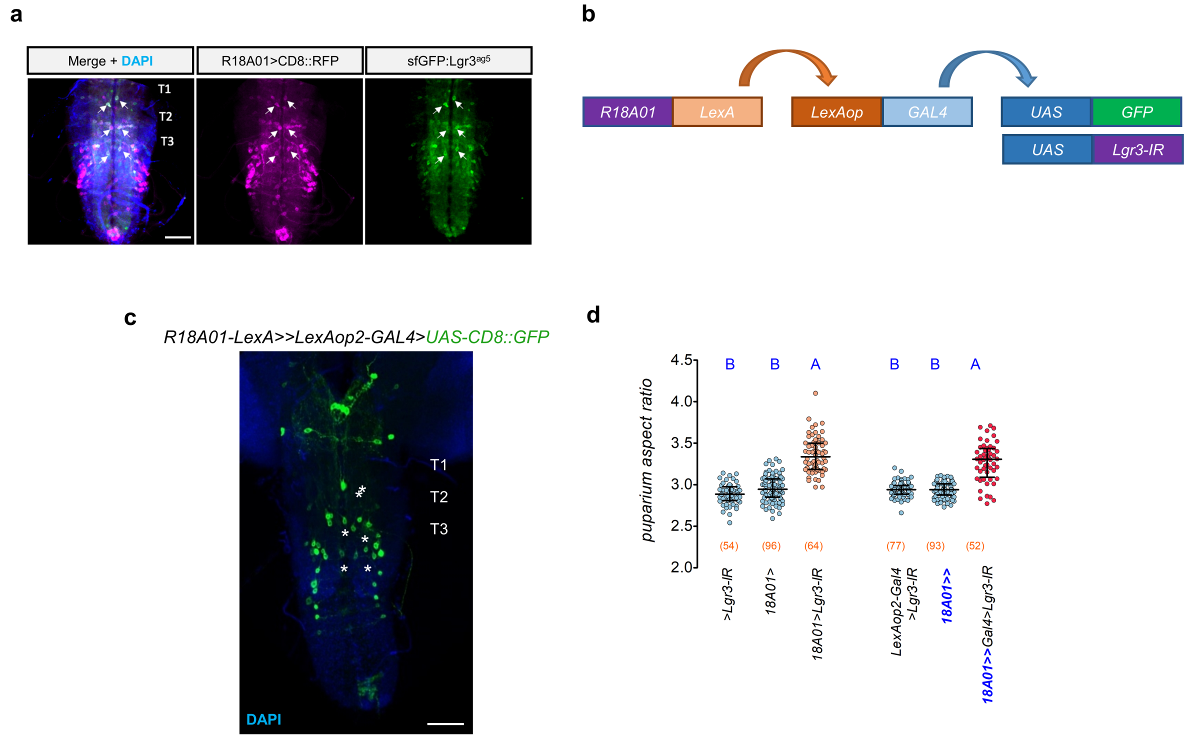


**Supplementary Fig. 9: *R18A01* expression pattern and validation of *R18A01-LexA* tools.** **a, c** Projections of confocal sections of a dissected white prepupa T0 CNS. **a** Six thoracic (6VNC) interneurons (white arrows) co-express *R18A01>CD8::RFP* (magenta) and *sfGFP::Lgr3$^{ag5}$* (anti-GFP, green). DAPI, blue. **b** Strategy for testing whether or not the *R18A01-LexA (R18A01>>)* line drives expression in the same *Lgr3*-sensitive cells that are important for pupariation behavior control as the *R18A01-GAL4* (*R18A01>*) line. *R18A01>>* drives expression of LexA which drives expression of GAL4 (*LexAop-GAL4*, *13xLexAop2-GAL4v-VP48*), which then drives expression of either *UAS-Lgr3-IR* or *UAS-CD8::GFP*. This is critical to generate a functional intersection, such as the *R18A01∩R48H10* intersectional genetics system. **c** *R18A01>>CD8::GFP* is expressed in the same 6VNC cells (asterisks) as *R18A01>*. T1-3, thoracic segments. **d** Dot plots of puparium aspect ratio (AR). *Lgr3* knockdown under the control of *R18A01>* or *R18A01>>* increases puparium AR. Statistics: **d** Horizontal bar, median. Error bars, 25-75%. Same blue letters, $P>0.05$. Dunn's test. (N) Number of animals (red).

**Supplementary Fig. 10.**

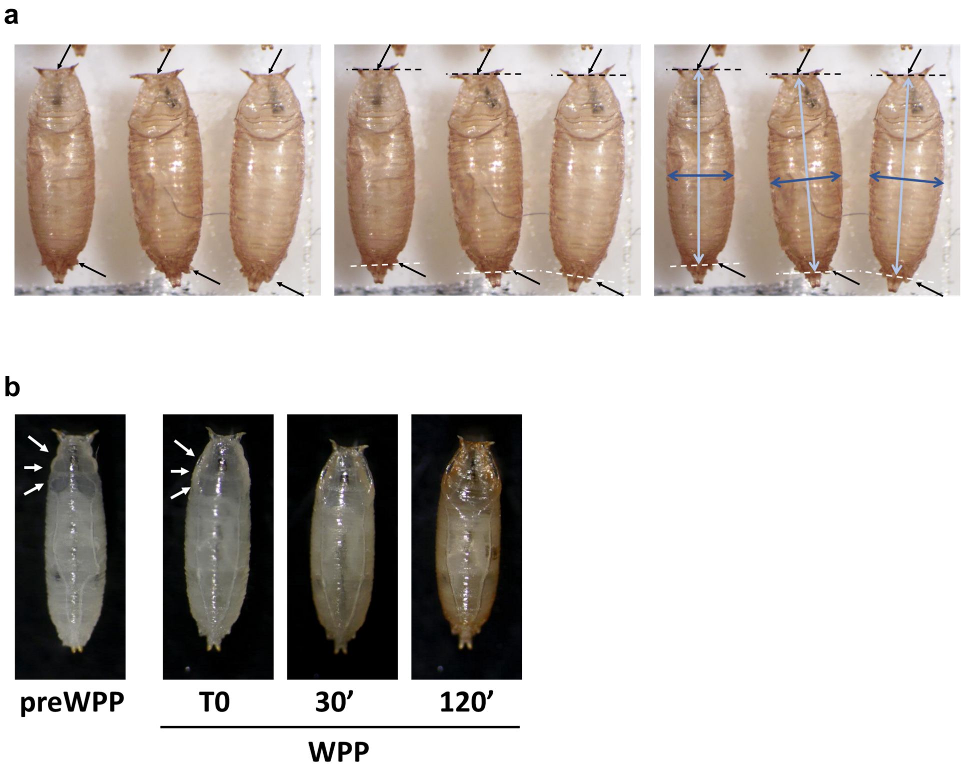


**Supplementary Fig. 10: Criteria for puparium aspect ratio and pupa staging**

**a** Measurement of the AR. The black arrows point to the anterior edge and the most anterior anal papillae that were taken as reference to measure the length of the pupa (light blue arrows). A perpendicular line drawn in the widest region of the pupae was used to measure the width. Aspect ratio was calculated as Length/Width. **b** Morphological features of a white prepupa (WPP = T0). A larva is considered to have reached the WPP stage when the operculum first becomes evident, the most anterior segments flatten in the dorso-ventral direction, and their edges thicken and lose the wiggly appearance (arrows). preWPP = animals that have started pupariation, but have not yet reached the WPP stage. Detectable tanning at WPP occurs ~30 min after T0.

**Supplementary Fig. 11.**

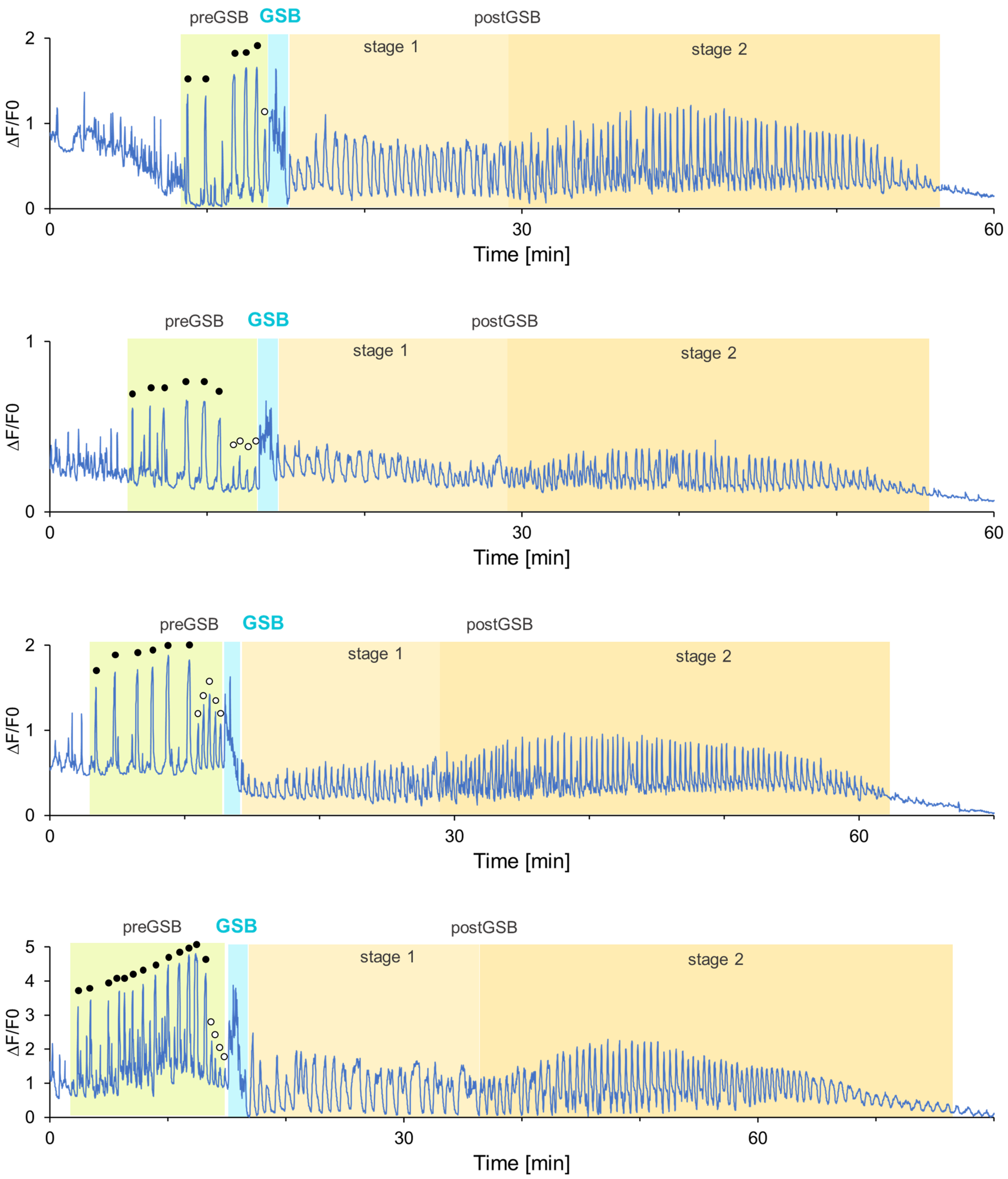


**Supplementary Fig. 11: Examples of mhc>>GCaMP fluorescence traces of WT animals in which the different components of the pupariation motor program (PMP) are depicted.**

The PMP begins once the larva has stopped wandering (see Fig. 4). Pre-GSB contractions are marked with a black circle. White circles indicate small fluctuations of mhc>>GCaMP signal intensity that occur between the last strong pre-GSB contraction and GSB and are the result of milder and shorter body contractions. $\Delta F/F0 = (F\text{-}F0)/F0$, being F0 the minimum value of each trace. Genotypes: Top three, *Lgr3*$^{ag2/ag1}$ *(+/-)*.. Bottom, *dilp8*$^{ag52/ag54}$ *(+/-)*.

### Supplementary Fig. 12.

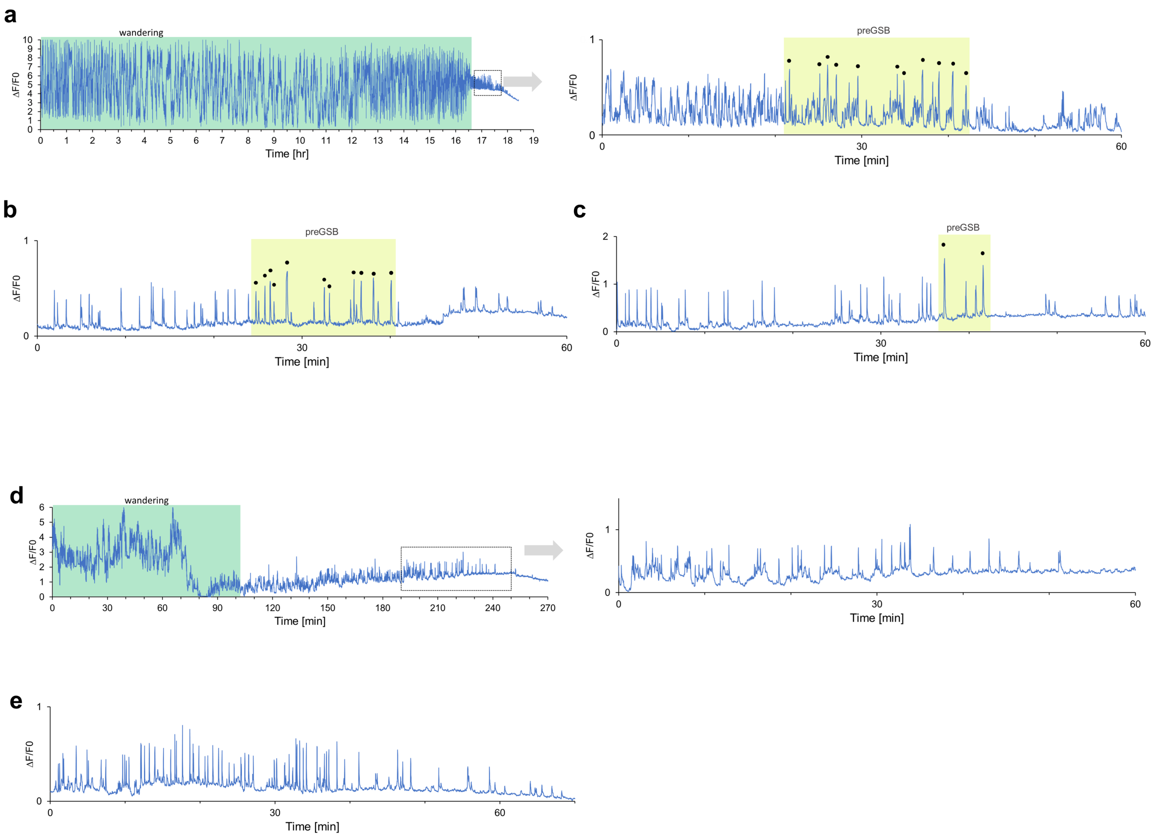


**Supplementary Fig. 12: Examples of mhc>>GCaMP-fluorescence traces of *Lgr3* mutant animals.**

**a** Similarly to WT animals, *Lgr3*$^{ag1/ag1}$ *(-/-)* mutant larvae wander along the arena for a variable time, sometimes as long as 16 h, until they stop and select a place to pupariate. Next, most animals perform pre-GSB-like contractions, but fail to progress further in the pupariation motor program. The graph on the right is an expansion of the region delimited by the dashed-line box, in which the peaks in total mhc>>GCaMP-fluorescence signals that result from whole body contractions that have been indicated with a black circle. **b, c** mhc>>GCaMP-fluorescence traces of two different mutant larvae that perform pre-GSB-like contractions, representative of the variability observed among individuals. **d, e** Examples of mutant larvae with no detectable whole-body preGSB contraction. ΔF/F0 = (F-F0)/F0, being F0 the minimum value of each trace.

### Supplementary Video Legends (1-10)

#### Supplementary Video 1.

Crawling wild-type larva expressing *mhc>>GCaMP* imaged under blue light under a dissecting microscope.

#### Supplementary Video 2.

Crawling wild-type larva expressing *mhc>>GCaMP* imaged under blue light in the pupariation-monitoring device used to track muscle contractions during pupariation.

#### Supplementary Video 3.

*mhc>>GCaMP* wild-type larva performing pre-GSB and GSB. The larva performs 5 pre-$\mathrm{GSB}^{\mathrm{short}}$ contractions and 4 pre-$\mathrm{GSB}^{\mathrm{long}}$ contractions, characterized by the activation of longitudinal ventral muscles that retract the anterior segments, before it performs GSB. The time of each contraction is indicated in the video.
The video is reproduced at 5X higher speed. Total elapsed time is 13 minutes.

#### Supplementary Video 4.

Wandering wild-type ($dilp8^{ag52/ag54}$) and mutant ($dilp8^{ag50/ag54}$) animals imaged under white light in the pupariation-monitoring device. Only the last pre-GSB contractions were included in the video.
The actual time elapsed at 10X speed is 10 minutes and at 60X speed is 2 h and 20 minutes.

#### Supplementary Video 5.

Sgs3::GFP wild-type larva expulsing the content of the salivary and spreading it over the ventral surface.

#### Supplementary Video 6.

*mhc>>GCaMP* and *Sgs3::GFP* wild-type larva performing pre-GSB and GSB. The larva performs three pre-GSB contractions (at 00:03, 00:15 and 00:32 min) before expulsing glue and progressing to GSB at 1:15 min.
The video is reproduced at 4X higher speed. Total elapsed time is 6.4 minutes.

#### Supplementary Video 7.

*mhc>>GCaMP* wild-type larva performing pre-GSB, GSB and part of post-GSB. The larva performs 8 pre-GSB contractions (at 0:15, 0:39, 01:02, 01:21, 01:38, 01:50, 02:03 and 02:11 min). Next, it enters in the GSB phase (from 02:26 to 02:42) and after that, it deploys post-GSB patterned contractions. Transient contraction of the muscles in the region that will form the operculum (>) can be observed from ~05:40 and become continuous around 7:30 min. The recording is interrupted for 1-2 minutes at 7:30. The video is reproduced at 4X higher speed. Total elapsed time is 32.6 minutes (2.5 min pre-GSB, 64 secs GSB and 29 min post-GSB). Tetanic contractions in the operculum occur ~20 minutes after GSB.

**Supplementary Video 8.**

*mhc>>GCaMP* wild-type larva performing pre-GSB, GSB and part of post-GSB. The larva performs 3 pre-GSB contractions (at 0:02, 0:06 and 00:14 min, the preceding contractions were not recorded). Then, it completes GSB (from 00:23 to 00:35) and begins postGSB contractions. Contractions that shape the operculum can be observed from ~04:45 and are labeled in the video with >.

The video is reproduced at 4X higher speed. Total elapsed time is 36 minutes. Contractions in the operculum occur ~17 minutes after GSB.

**Supplementary Video 9.**

*Drosophila virilis* wandering wild-type larva imaged under white light in the Pupariation monitoring device. They perform GSB at 00:36, 00:25 and 00:34 (left, center, right respectively).

The video is reproduced a speed 10X the real one. Total elapsed time is 10 min.

**Supplementary Video 10.**

*tub-gal80*$^{ts}$ *tub>dilp8 dilp8*$^{(-/-)}$ larvae developed at 18C express *dilp8* when shifted to 30 °C and perform GSB twice. After the first GSB (at 00:14 min), the larva resumes wandering and moves to a different place where it performs a second GSB (at 01:56 min). Tanning becomes evident at ~03:15 min.

The video is reproduced at 3X and 60X higher speed. The total time elapsed is 2 hours and the time between the two sequential GSBs is 39min 11sec.
